# Links between autobiographical memory richness and temporal discounting in older adults

**DOI:** 10.1101/765537

**Authors:** Karolina M. Lempert, Kameron A. MacNear, David A. Wolk, Joseph W. Kable

## Abstract

When making choices between smaller, sooner rewards and larger, later ones, people tend to discount or devalue future outcomes. This propensity can be maladaptive, especially as individuals age and their decisions about health, investments, and relationships become increasingly consequential. Individual differences in temporal discounting in older adults have been associated with episodic memory abilities, as well as with cortical thickness in the entorhinal cortex. The mechanism by which better memory might lead to more future-oriented choice remains unclear, however. Here we used a fine-grained measure of autobiographical memory richness (Autobiographical Interview scoring protocol) to examine which categories of episodic details are associated with temporal discounting in cognitively normal older adults. We also examined whether recalling autobiographical memories prior to choice can alter temporal discounting in this group. Time, place, and perceptual details, but not event or emotion/thought details, were associated with temporal discounting. Furthermore, time, place, and perceptual details and temporal discount rates were associated with entorhinal cortical thickness. Retrieving autobiographical memories prior to choice did not affect temporal discounting overall, but the extent to which the memories were rich in event and time details predicted whether they would reduce discounting after they were recalled. Thus, more future-oriented decision-makers may have more contextual (i.e., time, place, and perceptual) details in their recollections overall, and retrieving central event details at the time of choice may shift decisions toward being more patient. These findings will help with the development of interventions to nudge intertemporal choices, especially in older adults with memory decline.

---

Throughout our lives, we face many intertemporal choices that involve trade-offs between smaller, immediate gains and larger, long-term benefits (Strotz, 1956). For example, an individual may have to decide whether or not it is worth paying a fee to withdraw retirement funds early. People generally tend to prefer immediate rewards to delayed ones, and they discount the value of delayed rewards to a greater extent as the delay to receiving them increases. This tendency towards temporal discounting is nearly universal, but the rate at which people discount future rewards varies widely (Peters & Büchel, 2011). Steep temporal discounting, or overvaluing the present at one’s long-term expense, is associated with risky behaviors such as alcohol use (Vuchinich & Simpson, 1998), gambling (Reynolds, 2006), smoking (Bickel, Yi, Kowal, & Gatchalian, 2008; Yi & Landes, 2012), and excessive credit card borrowing (Meier & Sprenger, 2010). Given the substantial negative impact of steep temporal discounting, it is important to understand the neurocognitive mechanisms that underlie intertemporal choices, and to use this knowledge to develop interventions that could reduce impatience. Understanding how changes in cognition in healthy aging may influence temporal discounting is especially important, as older adults make many consequential intertemporal choices (e.g., about investments and health care).

One neurocognitive system that may support future-oriented intertemporal choices, and that declines with aging (Buckner, 2004), is episodic memory, or context-rich memory for autobiographical events (Tulving, 1987). It has been proposed that the ability to retrieve detailed episodic memories could enable a more vivid imagination of the future, which would then enhance the value of future rewards (Boyer, 2008; Trope & Liberman, 2000). In line with this, we recently found that in older adults, better memory abilities (specifically, better episodic memory recall and semantic fluency) were associated with lower temporal discounting rates (Lempert et al., 2019). Structural integrity in the medial temporal lobe (specifically, in the entorhinal cortex) was also associated with temporal discounting in this group (Lempert et al., 2019). A limitation of this previous study, however, is that it examined only standard neuropsychological measures of memory, which may not fully capture age-related decline in episodic memory in cognitively normal older adults. With these measures, we were also unable to establish whether semantic or episodic memory has a greater impact on temporal discounting. Therefore, it remains unclear whether detailed episodic memory is critical for making more patient choices.

Here we use a more fine-grained measure of episodic memory – autobiographical memory richness – to examine which aspects of autobiographical memory are associated with temporal discounting. Not only might autobiographical memory retrieval be more directly relevant to intertemporal decision-making, but the quality of autobiographical memories changes over the lifespan, becoming more schematic in healthy older adults (Levine, Svoboda, Hay, Winocur, & Moscovitch, 2002). So far, studies that have investigated the link between autobiographical memory richness and temporal discounting have yielded null results (Bromberg, Wiehler, & Peters, 2015; Seinstra, Grzymek, & Kalenscher, 2015). It is possible, however, that these studies failed to detect an association because they used a *composite* measure of episodic details. The Autobiographical Interview scoring protocol (Levine et al., 2002) and other similar protocols (Irish et al., 2011) distinguish among different categories of details in autobiographical memories, such as central event details and peripheral perceptual details. These different categories might differentially relate to intertemporal decision-making. For example, while time, place, and perceptual details are considered *contextual* and reflect mental construction of a specific scene, event and emotion details are considered *story* details that may reflect an individual’s retelling of a schema or script. Here, in a group of older adults who described autobiographical memories, we examined the relationship between temporal discounting and five different categories of episodic details: time, place, perceptual, event, and emotion/thought details.

There is some neural evidence consistent with the idea that different categories of details tap into different aspects of episodic memory retrieval. For example, individuals with unilateral temporal lobe epilepsy resulting in damage to the hippocampus show deficits in time, place, perceptual, and emotion details, but not event details, suggesting that story elements can remain intact even with hippocampal damage (St-Laurent, Moscovitch, Levine, & McAndrews, 2009). Moreover, different forms of dementia result in different profiles of autobiographical memory recall. Individuals with semantic dementia show impairments in spatiotemporal details, but spared event details, whereas individuals with Alzheimer’s disease show impaired event details, with preserved perceptual and spatiotemporal details (Irish et al., 2011). A subset of participants here had structural MRI data, allowing us to segment the medial temporal lobe to examine which subregions are associated with different categories of autobiographical details, as well as with temporal discounting. Given that entorhinal cortical thickness was associated with temporal discounting in a previous study (Lempert et al., 2019), we were especially interested in interrelationships between entorhinal cortex, autobiographical details, and temporal discounting in this sample.

In addition to correlational evidence linking temporal discounting with episodic memory ability, previous research also shows that the most effective *manipulations* of temporal discounting involve engaging the episodic memory system. Imagining positive future events decreases temporal discounting in young adults (Benoit, Gilbert, & Burgess, 2011; Palombo, Keane, & Verfaellie, 2015; Peters & Büchel, 2010; Sasse, Peters, Büchel, & Brassen, 2015), whether or not the future events are directly linked to the future rewards at stake (Peters & Büchel, 2010). Recently, positive autobiographical memory retrieval has also been shown to reduce discounting in young adults (Lempert, Speer, Delgado, & Phelps, 2017). Just as it is unknown which aspects of episodic memory are associated with temporal discounting across individuals, the mechanism by which memory-based manipulations increase patience also remains unclear. Previous research suggests that more vivid episodic imagery would lead to more patient choice (Peters & Büchel, 2010), but previous studies have examined only self-reported ratings of vividness. Here we took advantage of our more objective autobiographical memory scoring analysis to investigate (1) whether recalling these autobiographical memories would influence intertemporal choice in older adults, and (2) which autobiographical details, if any, predict a change in temporal discounting after memory recall. We hypothesized that, as a group, older adults would show a blunted effect of memory recall on choice, due to their memory decline. Scoring descriptions of these memories, however, allowed us to investigate the extent to which any effect of the manipulation can be explained by inter-individual variability in autobiographical details, or intra-individual (i.e., between memories) variability in autobiographical details.

In sum, this study aims to determine the following in a group of cognitively normal older adults: (1) the relationship between temporal discounting and autobiographical memory richness, (2) the relationship between medial temporal lobe structural integrity and both temporal discounting and autobiographical memory richness, and (3) whether positive autobiographical memory recall impacts temporal discounting in this group.

## Method

### Participants

Thirty-eight older adult participants (ages 65-90; mean age = 74; SD = 6.9; 24 F, 14 M; 31 White, 6 Black, 1 Asian) completed this experiment. All subjects were deemed cognitively normal based on consensus diagnosis at the Penn Alzheimer’s Disease Core Center. Of the thirty-eight participants who completed the study, four were excluded. Three were excluded because their discount rates could not be estimated in one or both experimental conditions. Of these 3, two chose all delayed rewards, and 1 chose all immediate rewards. An additional participant was excluded for having a score in the moderately depressed range on the Geriatric Depression Scale. Thus, thirty-four participants were included in final analyses (23 F; mean age = 74.11; SD = 6.97). All participants were compensated $10/hour for their participation. The study was approved by the Institutional Review Board of the University of Pennsylvania.

### Procedure

Participants completed a two-day study. On Day 1, they described positive memories prompted by each of 12 life event cues (e.g., “being in a wedding,” “winning an award”). The cues were a compilation of cues from prior studies (Sharot, Riccardi, Raio, & Phelps, 2007; Speer, Bhanji, & Delgado, 2014) and were designed to probe for positive memories. For each cue, participants selected a memory in which they had been personally involved and that had occurred at a specific place and time. For each memory, participants had four minutes to provide a brief verbal description, and their response was audio recorded for later scoring with the Autobiographical Interview Protocol (Levine et al., 2002; see *Autobiographical memory scoring* section below). They were prompted when there was one minute left. Each memory cue was allotted four minutes, even if participants did not speak through the whole four minutes. An interviewer was present throughout this task, in order to prompt the participant for more detail if necessary. At the end of each memory description, they provided the location and date of the memory. Then, they gave subjective ratings for valence (1 = neutral; 2 = positive), emotional intensity (1-4: 1 = not intense, 4 = very intense), feeling now (i.e., how they felt when recalling the memory; 1-4: 1 = neutral, 4 = very good), feeling at the time of the memory (1-4: 1 = neutral, 4 = very good), personal importance of the memory (1-4: 1 = not important; 4 = extremely important), similarity between current self and self in the memory (1-4: 1 = very different; 4 = exactly the same) and vividness (1-4: 1 = not vivid, 4 = very vivid). A table with average memory ratings is presented in Supplemental Table 1 in the online supplemental materials. Participants were instructed to select memories that were positive. If they could not think of a specific memory for a cue, or if the cue was only associated with negative memories, they could skip the cue and receive another. There were 24 possible cues, but the experiment terminated after twelve memories had been described (see Supplemental Methods in online supplemental materials for list of cues). This procedure was adapted from a previous study of episodic recall (Bertossi, Tesini, Cappelli, & Ciaramelli, 2016).

In preparation for the second session, nine of each participant’s positive memories were selected. These nine had been rated as positive (i.e., valence = 2), and had the highest combined intensity and feeling ratings. They were summarized in subject-specific event cues that the participants reviewed at the beginning of the second session, to ensure that they could bring to mind the memory associated with each cue.

Participants returned for the second session about one week later (M = 7.15 days; SD = 1.85; range: 2-14 days) to perform an intertemporal choice task. On each trial of this task, they were presented with a screen showing two options: “$10 today” and a monetary reward of larger magnitude available after a delay (e.g., “$20 in 30 days”). Delayed reward amounts varied from $11 to $35, and delays varied from 1 day to 180 days (see Supplemental Methods in online supplemental materials for list of all amounts and delays). An effort was made to capture a range of hyperbolic discount rates (range: 0.00018 – 0.25) with the constraints that the immediate amount always be $10 and the delay not exceed 180 days. The immediate reward was kept constant, so that this paradigm closely resembled the previous version of this paradigm used in younger adults (Lempert et al., 2017). For each choice, participants made a button press, indicating which option they preferred (the task was self-paced). The order of the trials was randomized, and the immediate and delayed reward options switched sides of the screen randomly. After participants responded, they were shown the option they had just chosen for 1 second. After a 2 second inter-trial interval, the next choice screen appeared. There were 54 distinct choices, shown once in each condition, for a total of 108 trials.

Participants made these choices in two conditions, “Memory” and “Control,” presented in blocks. In Memory blocks, participants re-accessed the nine positive memories (yielding nine “mini-blocks”) triggered by cues from Day 1 before making choices. At the beginning of each Memory mini-block, a fixation point appeared for 3 seconds. Then, a memory cue was displayed for 20 seconds. Participants were asked to recall the memory described by this cue and to elaborate on it for as long as they could or until 20 s were up. After a 3 s inter-stimulus interval, participants rated the memory on valence, emotional intensity, and feeling (allotted 4 s for each). Following this, participants made 6 intertemporal choices before the next memory cue appeared on the screen. The first memory block consisted of 5 mini-blocks (5 memories and 30 intertemporal choices), and the second memory block consisted of 4 mini-blocks (4 memories and 24 intertemporal choices; Fig. 1).

**Fig. 1.**
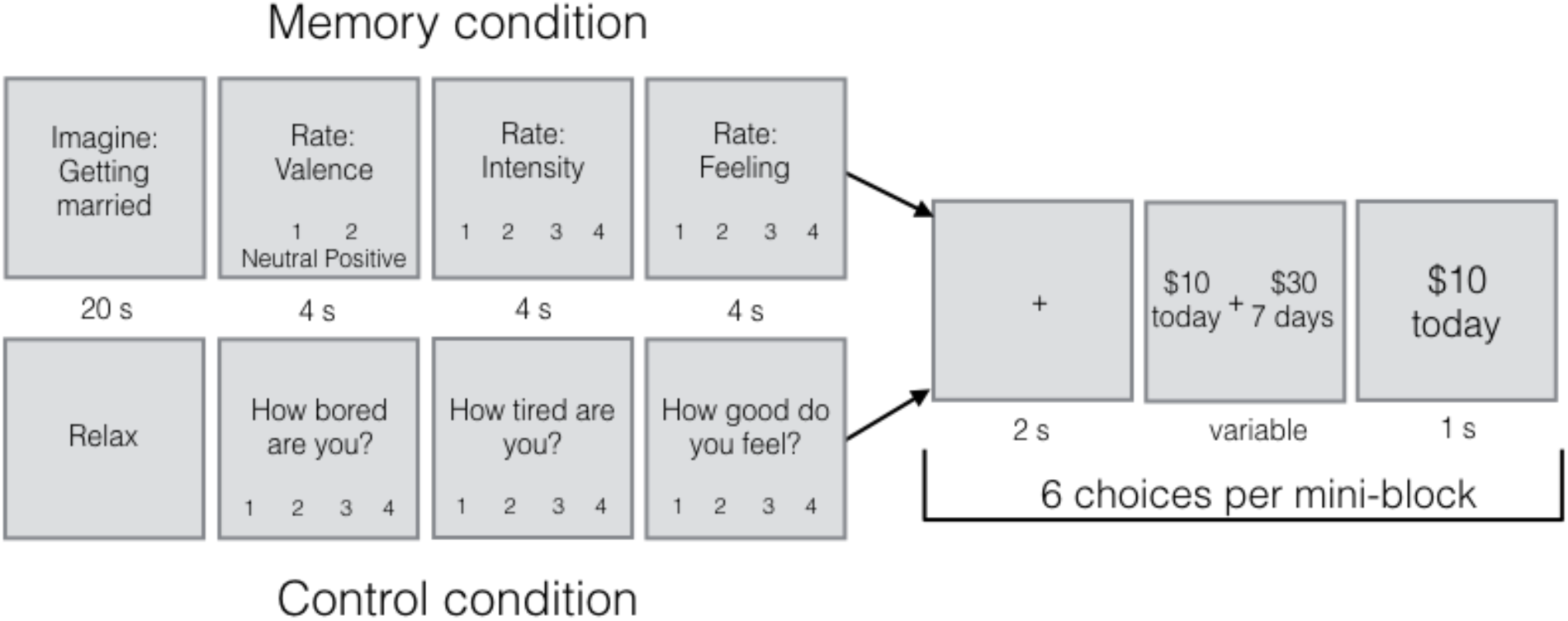
Task Layout. Each mini-block contained six intertemporal choices. Each memory mini-block began with a memory cue, describing an autobiographical memory specific to the participant. The participant was asked to think about that positive memory for 20 seconds. Then they rated the valence (1 = neutral; 2 = positive), intensity (1-4; 1 = not intense; 4 = very intense) and feeling (1-4; 1 = neutral; 4 = very good) of the memory. Finally, they made 6 choices between $10 today and a larger amount of money available after a delay. The participant made a button press while the options were on the screen, and then was shown what they chose for 1 s before the next trial began. In the Control mini-blocks, participants were told to relax for 20 s and then to answer questions about how bored and tired they were, and how good they felt (1-4 scale for each). They then made the same intertemporal choices in this condition. This procedure is adapted from (Lempert et al., 2017).

In each of the Control mini-blocks, participants first saw the word “Relax” on the screen for 20 s. They were instructed to rest during this time. Then, they rated how tired they were (1-4; 1 = very awake; 4 = very tired), how bored they were (1-4; 1 = not bored; 4 = very bored), and how good they felt (1-4; 1 = neither good nor bad; 4 = very good; 4 s for each rating). Following this, they made 6 intertemporal choices before the next “relax” screen appeared. The first Control block consisted of 5 mini-blocks (5 “relax” screens and 30 intertemporal choices), and the second Control block consisted of 4 mini-blocks (4 “relax” screens and 24 intertemporal choices). There were two Control blocks and two Memory blocks, which alternated; the block type that came first was counterbalanced across subjects. The same choices were presented in both conditions. The task was programmed using E-Prime 2.0 Stimulus Presentation Software (Psychology Software Tools, Sharpsburg, PA).

Participants were told at the outset of the Day 2 session that one of the intertemporal choice trials would be randomly selected and they would receive the amount they chose on that trial, at the delay specified. They were paid via a pre-paid debit card (Greenphire Clincard system). If they chose the immediate reward on that trial, they would receive the money on their card that day. If they chose the delayed reward, they would receive the money after the delay had elapsed. Paying out both reward types this way ensured that transaction costs were equivalent between immediate and delayed rewards.

After the decision making task was completed, participants filled out four questionnaires on a computer: the Interpersonal Reactivity Index (IRI; Davis, 1983), the Life Orientation Test-Revised (LOT-R; Scheier, Carver, & Bridges, 1994), the Geriatric Depression Scale (GDS; Yesavage, 1988), and the Vividness of Visual Imagery questionnaire (VVIQ; Marks, 1973). The GDS was included as a screening tool, because symptoms of depression are associated with deficits in memory ability, especially in positive memory recall (Dillon, 2015). Therefore, anyone with a GDS score of 9 or above (out of 15), indicating moderate or severe depression, was excluded (n = 1; see *Participants* above). Details of and results from analyses of these questionnaires can be found in Supplemental Methods, Supplemental Results, and Supplemental Fig. 1 in online supplemental materials.

### Analyses

#### Choice data

Participants’ individual intertemporal choice data were fit separately for choices in the Memory blocks and Control blocks with the following logistic function using maximum likelihood estimation:

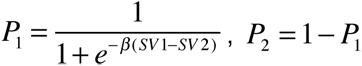

Here, P_1_ refers to the probability of choosing the delayed option, and SV_1_ and SV_2_ are the subjective values of the delayed and immediate options, respectively. The subjective value of the options was assumed to follow a hyperbolic discounting function (Kable & Glimcher, 2007; Mazur, 1987):

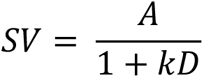

Where SV is the subjective value, A is the amount, D is the delay to receiving the reward, and *k* is the subject-specific discount rate parameter (higher *k* values correspond to more impatience). Since discount rates are not normally distributed, these parameters were log-transformed before statistical analyses were performed.

#### Autobiographical memory scoring

Audio recordings from each of the nine memories used in the intertemporal choice task for each participant in the final sample (N = 35) were transcribed and then scored using the Autobiographical Interview protocol (Levine et al., 2002). The transcription that was scored for each memory began after the participant selected the cue, and ended after the participant stated the date and location of the memory.

For each memory transcription, the central event was identified. If more than one event was mentioned, the event described in more detail (the same one also described by the cue on Day 2) was considered the main event. Each event was divided into distinct details (unique pieces of information), and these details were classified as internal to the event (episodic details) or external (information related to events other than the main event). Semantic information and repetitions were considered external. Internal details and external details that were not semantic details or repetitions were further categorized as: event details (describing happenings in the story), time details (pertaining to when the event occurred), place details (pertaining to where the event occurred), perceptual details (describing sensations during the event), and emotion/thought details (pertaining to the mental state of the subject at the time of the event).

One rater who was blind to the hypothesis of the study scored the transcripts. An additional rater scored a subset of memories (approximately 45% of all memories; 142 from 16 participants) independently, to assess inter-rater reliability. Across all memories scored by both raters, there was a high correlation between the total number of internal details (Pearson r = 0.74; p < 0.001), and the total number of external details (r = 0.76; p < 0.001).

#### Autobiographical details and individual differences in discount rate

Scoring autobiographical details allowed us to examine the relationship between autobiographical memory richness and temporal discounting across participants. We averaged the number of internal details across all nine memories for each participant, in each category (event, time, place, emotion/thought, and perceptual). Spearman correlations were conducted between the Control condition discount rate and (1) average internal event details, (2) average internal time details, (3) average internal place details, (4) average internal emotion/thought details, and (5) average internal perceptual details.

In addition, we explored the relationship between age and temporal discounting, as well as between age and different categories of autobiographical details. We expected to find no relationship between age and temporal discounting, consistent with previous work (Lempert et al., 2019).

#### Structural MRI data acquisition and analysis

Twenty-two participants in the sample also underwent MRI scanning within approximately one year of participating in this experiment (mean number of days between MRI and first behavioral testing session = 213.55; SD = 112; range: [14, 405]). MRI data were obtained on a Siemens Prisma 3T MRI with a 64-channel head coil. T1-weighted high-resolution magnetization-prepared rapid-acquisition gradient echo (MPRAGE; 0.8 x 0.8 x 0.8 mm^3^ voxels; TR/TE/TI=1600/3.87/950 ms; flip angle=15°) anatomical scans were collected. The medial temporal lobe was segmented using an automatic pipeline, ASHS–T1 (Xie et al., 2019). This technique uses a multi-atlas label fusion approach (Wang et al., 2013) together with a tailored segmentation protocol to take into account anatomical variability in MTL cortex. It also explicitly labels dura, which has similar appearance to gray matter in T1-weighted MRI, resulting in more accurate segmentation of MTL cortex compared to other T1-MRI segmentation pipelines, such as FreeSurfer (Xie et al., 2019). In addition to a volume measure of the hippocampus, we obtained measures of mean cortical thickness in the following regions-of-interest (ROIs): entorhinal cortex (ERC), perirhinal cortex subregions BA35 and BA36, and parahippocampal cortex (PHC), using a graph-based multi-template thickness analysis pipeline (Xie et al., 2017). We conducted a series of exploratory regression analyses to investigate relationships between thickness/volume in each of these subregions and temporal discounting, as well as with each category of internal autobiographical details (event, time, place, emotion/thought, and perceptual). Thickness and volume measures were averaged across hemispheres. Image quality was inadequate for segmentation of ERC, BA35, and BA36 for one participant (leaving n = 21 for those analyses). For another participant, image quality was inadequate for BA36 on one side only, so the mean was replaced with mean cortical thickness from the available side. Age, gender, years of education, and the absolute number of days between the MRI scan and the first behavioral testing session were entered as covariates of no interest. For the hippocampal volume analysis, we included the (square root-transformed) intracranial volume as an additional covariate. Partial Pearson correlation coefficients are reported.

#### Positive autobiographical memory manipulation analysis

To test for the effect of our positive memory manipulation on temporal discounting overall, we conducted a two-tailed paired *t*-test to compare discount rates between Memory and Control conditions for each participant. We also investigated which, if any, details of the memories themselves predicted the extent to which they were effective in reducing discount rate. To this end, we constructed a mixed-effects logistic regression predicting choice (0 = chose $10 immediate reward; 1 = chose delayed reward) on all trials in the Memory condition. We conducted one regression for each of the five categories of internal details (event, time, place, emotion/thought, and perceptual). Since memory descriptions vary with respect to total verbal output, we looked at the *percentage* of internal event, time, place, emotion/thought and perceptual details relative to the total (internal + external) number of details in that category. In each regression, we controlled for the subjective value of the delayed option on that trial, assuming the participant’s discount rate from the Control condition. Specifically, we plugged in the discount rate *k* that was fitted to the data in the Control condition only, along with the amount and delay on that particular trial into the hyperbolic model equation, to determine the subjective value of the delayed reward on that trial, and entered this as a nuisance regressor. This is a conservative test that allowed us to see whether autobiographical details could predict delayed reward choice in the Memory condition *above and beyond* what could be predicted from the discount rate in the Control condition trials alone. We allowed slopes (for the regressor of interest) and intercepts to vary by subject. We conducted a similar analysis with the participants’ ratings of their memories as independent variables, yielding null results. The results of that analysis are presented in Supplemental Results in the online supplemental materials.

In addition to examining within-subject variance in the effect of memory on discounting, we also examined whether individual differences in autobiographical details (i.e., average internal event details, average internal time details, average internal place details, average internal emotion/thought details, and average internal perceptual details) or differences in age predicted the effect size of our manipulation (difference between log-transformed discount rate in Control condition and Memory condition) across participants.

## Results

### Individual differences in contextual autobiographical details are associated with temporal discounting

Temporal discounting was correlated with the number of contextual details in autobiographical memories. To understand which aspects of autobiographical memory, if any, were associated with temporal discounting, we examined associations between the average number of event, time, place, perceptual, and emotion/thought details in each participant’s autobiographical memory descriptions and their temporal discounting rate. We found that the average number of internal event details (ρ = -0.08; p = 0.649) and emotion/thought details (ρ = - 0.13; p = 0.475) were not associated with discount rate, but internal time details (ρ = -0.46; p = 0.006), place details (ρ = -0.54; p < 0.001), and perceptual details were (ρ = -0.36; p = 0.039; Fig. 2). Note, however, that only the associations with time and place details survive correction for multiple comparisons (Bonferroni corrected α level = 0.01). Furthermore, after regressing out variance in discount rate due to age, gender, and years of education, the association remained significant for time details (ρ = -0.43; p = 0.011) and place details (ρ = -0.43; p = 0.012), but not for perceptual details (ρ = -0.25; p = 0.150). Unlike event and emotion details, time, place, and perceptual details are considered *contextual* autobiographical details, which localize memory to a particular place and time. This specificity in the kinds of details related to temporal discounting suggests that our results are not driven by individual differences in verbal output alone. To further control for overall verbal output, however, we ran a correlation between temporal discounting and the average number of *external* (non-episodic) details in participants’ autobiographical memory descriptions and found no relationship (ρ = -0.002; p = 0.990).

**Fig. 2.**
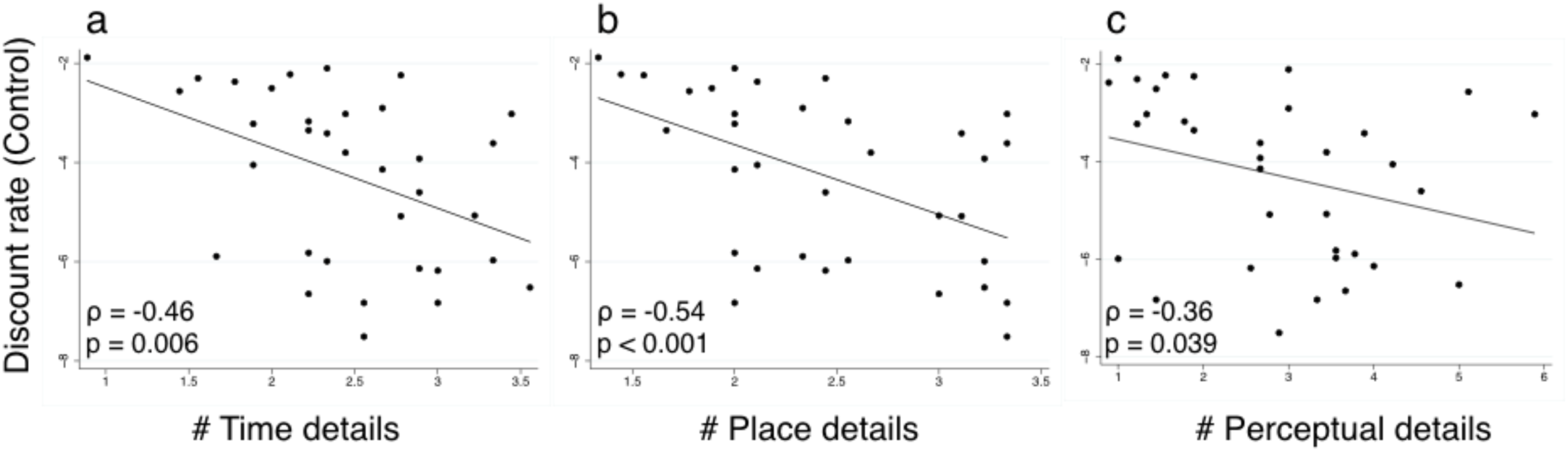
Association between average number of internal time (a), place (b), and perceptual (c) details across nine autobiographical memories and temporal discounting rate. Temporal discounting rate in the Control condition of the intertemporal choice task was used as the dependent variable. Individuals who mentioned more time, place, and perceptual details were more likely to select larger, later rewards in the temporal discounting task.

Neither temporal discounting nor time and place details were associated with age. We found no relationship between age and temporal discounting (ρ = 0.19; p = 0.293), or between age and internal event (ρ = -0.19; p = 0.287), time (ρ = -0.12; p = 0.483), or place (ρ = -0.13; p = 0.449) details. However, perceptual (ρ = -0.45; p = 0.008) and emotion/thought details (ρ = - 0.42; p = 0.014) were associated with age, such that older individuals mentioned fewer perceptual and emotion/thought details in their memory descriptions.

### Entorhinal cortical thickness is associated with temporal discounting and contextual autobiographical details

Given that our previous research found that temporal discounting was associated with structural integrity in ERC in older adults (Lempert et al., 2019), we also examined this relationship in the subset (n = 22) of participants who had structural MRI data, which partially overlapped with our previous study. Consistent with our previous results, mean ERC thickness was associated with temporal discounting (r = -0.63; p = 0.007; Fig. 3), but thickness in BA35 (r = 0.09; p = 0.727), BA36 (r = 0.14; p = 0.585), PHC (r = -0.33; p = 0.177), and hippocampal volume (r = -0.32; p = 0.208) were not.

**Fig. 3.**
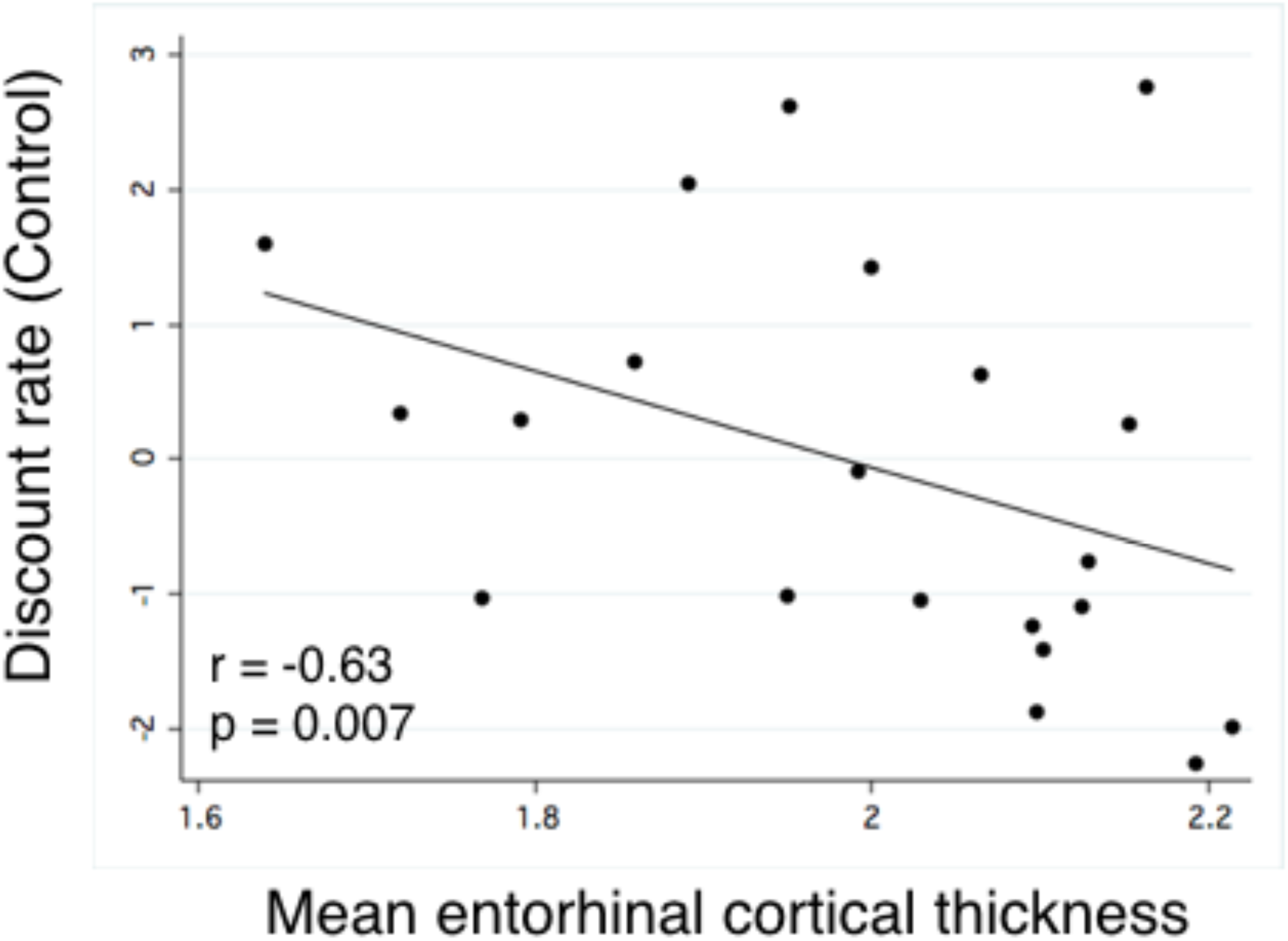
Individuals with more cortical thickness in entorhinal cortex displayed reduced temporal discounting, or a relatively greater preference for larger, delayed rewards (n = 21).

A follow-up exploratory analysis found that ERC thickness was most strongly associated with the autobiographical detail categories – place, time, and perceptual details – that are also associated with temporal discounting. We ran a series of regressions investigating relationships between categories of autobiographical details and ERC thickness. ERC thickness was significantly associated with the average number of place details (r = 0.71; p = 0.002), time details (r = 0.58; p = 0.014) and perceptual details (r = 0.62; p = 0.008), but not event (r = 0.26; p = 0.322), or emotion/thought details (r = 0.37; p = 0.143). The full list of partial correlation coefficients between autobiographical details and medial temporal lobe subregions is presented in Table 1. Other correlations of note are that hippocampal volume is significantly (p < 0.05) associated with time details, and PHC thickness is significantly associated with perceptual details.

**Table 1.** Associations between cortical thickness/volume in medial temporal lobe subregions and the average number of autobiographical details in each category.

| Medial temporal lobe region | Event | Time | Place | Perceptual | Emotion/<br>Thought |
| --- | --- | --- | --- | --- | --- |
| Entorhinal cortex | 0.26 | **0.58\*** | **0.71\*\*** | **0.62\*\*** | 0.37 |
| BA35 | 0.37 | 0.10 | 0.05 | -0.04 | 0.33 |
| BA36 | -0.02 | 0.25 | 0.24 | 0.01 | -0.15 |
| Parahippocampal cortex | 0.40 | 0.45 | 0.19 | **0.50\*** | 0.46 |
| Hippocampus (volume) | 0.34 | **0.59\*** | 0.10 | 0.18 | -0.17 |

Note: Partial Pearson correlation coefficients, controlling for age, gender, years of education and absolute number of days since MRI are shown. \* $p < 0.05$ , \*\* $p < 0.01$ (uncorrected for multiple comparisons).

### No effect of positive memory retrieval on temporal discounting in older adults

On Day 2, participants made a series of intertemporal choices either after retrieving positive autobiographical memories for 20 seconds (Memory blocks) or after relaxing for the same amount of time (Control blocks). Unlike in younger adults in a previous study (Lempert et al., 2017), the positive memory manipulation had no effect on temporal discounting in this older adult population (two-tailed t_33_ = -0.21; *p* = 0.834; Fig. 4). Surprisingly, on average, positive feeling ratings were not higher in the positive memory blocks compared to the control blocks (t_33_ = 1.09; p = 0.285), most likely because feeling ratings were close to ceiling in both conditions (mean feeling rating in Control blocks = 3.53; mean feeling rating in Memory blocks = 3.61; maximum feeling rating = 4).

**Fig. 4.**
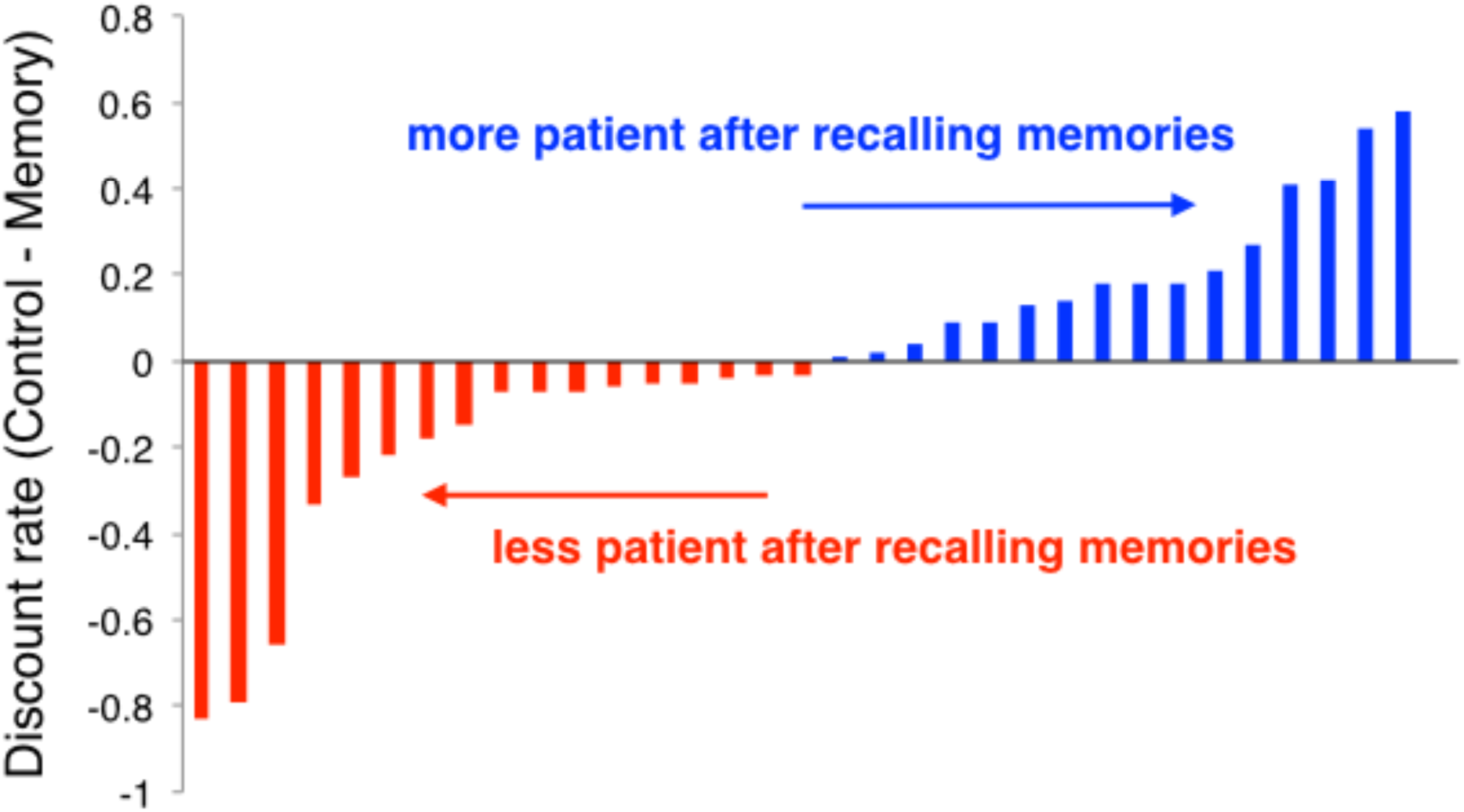
Positive memory retrieval does not affect intertemporal choice in older adults. The difference between the log-transformed discount rate in the Control condition and Memory condition is plotted for each subject. Positive difference (blue) indicates more patience in the positive memory condition. Negative difference (red) indicates more impulsivity in the positive memory condition. (t_33_ = -0.21; p = 0.834; N = 34; N = 1 not shown in figure because difference in discount rate was 0).

### Event and time details predict patient choice following memory recall within participants

Though recalling positive autobiographical memories did not have an overall effect on temporal discounting, individual memories with a high percentage of time and event details did bias choices towards delayed rewards. To investigate whether any aspects of the memories themselves could predict the extent to which retrieving them was effective in reducing temporal discounting rate, we conducted a series of mixed-effects logistic regressions to see which categories of internal memory details (if any) could predict choice of delayed reward on a trial-by-trial basis. Importantly, by entering the Control condition subjective value as a regressor in the model, this analysis controlled for the amount and delay offered on that trial, as well as the subject’s idiosyncratic discount rate in the Control condition. We found that memories that were more detailed in their description of what occurred (i.e., percentage of event details) were more likely to lead to delayed reward choice after they were recalled (Coefficient = 1.114; p = 0.033). Memories that contained a higher percentage of internal time details (Coefficient = 0.855; p = 0.029) were also more likely to lead to more patient choice. Place (Coefficient = 0.052; p = 0.891), emotion/thought (Coefficient = 0.493; p = 0.271), and perceptual details (Coefficient = 0.476; p = 0.175) were not significant predictors of choice.

Although individual differences in time, place, and perceptual autobiographical memory details predicted temporal discounting rate at baseline, they were unrelated to the effect of memory retrieval on temporal discounting across individuals. That is, the difference in discount rate between Control and Memory conditions was not associated with internal event details (ρ = -0.17; p = 0.323), time details (ρ = -0.12; p = 0.509), place details (ρ = -0.24; p = 0.168), emotion/thought details (ρ = 0.09; p = 0.598), or perceptual details (ρ = -0.04; p = 0.821). Moreover, participant age did not significantly predict the manipulation effect size (ρ = -0.15; p = 0.390).

## Discussion

In the current study, we examined associations between autobiographical memory richness and temporal discounting in a group of cognitively normal older adults. We found that the average number of time, place, and perceptual details, but not event or emotion/thought details, in participants’ positive autobiographical memories was associated with individual differences in temporal discounting. In other words, people who were better able to localize their memories to a particular time and place, and who reported more sensations from that time and place, were more likely to choose larger, later rewards. Cortical thickness in entorhinal cortex was associated with these same contextual memory details as well as temporal discounting, suggesting that it may be a neural substrate connecting memory retrieval with decision-making. We also tested whether recalling these positive autobiographical memories would reduce temporal discounting in older adults. Although recalling positive memories reduces temporal discounting in young adults (Lempert et al., 2017), it was not effective in older adults. However, within participants, the extent to which autobiographical memories contained a greater proportion of episodic time and event details predicted whether choices became more future-oriented following recall of those memories.

To our knowledge, this is the first study to detect an association between autobiographical memory details and temporal discounting. Previous studies found no association between overall autobiographical memory richness and discount rate (Bromberg et al., 2015; Seinstra et al., 2015). By using a composite measure of internal (episodic) details, though, these studies may have masked the finding that discounting may be related to only select types of details. Here we found that time, place, and perceptual details, but not emotion/thought or event details, predicted more future-oriented decision-making. This suggests the intriguing possibility that people who are more future-oriented do not necessarily have more detailed memories in general, but rather, they are more likely to encode and/or recall particular kinds of details, such as those pertaining to the time and place of an episode. Time, place, and perceptual details are considered contextual details and thus may reflect the mental construction of a specific scene. It is worthy of mention that these contextual details are the same ones that are relatively more impaired in individuals with semantic dementia (Irish et al., 2011), who also show increased temporal discounting (Chiong et al., 2016) and impoverished episodic future thinking (Irish, Addis, Hodges, & Piguet, 2012) relative to healthy controls and people with Alzheimer’s disease.

In the subset of participants who had structural neuroimaging data, we replicated the finding from our previous study (Lempert et al., 2019) that thickness in ERC (and no other subregion of the MTL) was associated with temporal discounting rate. This replication is notable since the current study included only cognitively normal participants, who performed a different intertemporal choice task from the previous study. However, these results should be interpreted with caution, as this is a small sample (n = 21) that is partially overlapping with the prior study. In an exploratory analysis examining associations between autobiographical details and MTL structural integrity, we found that the same detail categories that were associated with temporal discounting – time, place and perceptual details – were the ones most closely associated with ERC thickness. This is the first study to our knowledge to link spatiotemporal details with ERC thickness, although this connection is consistent with the well-known role of ERC in spatial navigation and memory (Igarashi, 2016; Sugar & Moser, 2019). To date, the literature on autobiographical memory has focused primarily on the distinction between external and internal details, but our behavioral and neural results highlight the importance of considering different categories of internal details.

Here we also tested if recalling positive memories directly prior to intertemporal choice would alter temporal discounting in older adults, given that they show a marked decline in episodic memory (Koen & Yonelinas, 2016; Levine et al., 2002; Nyberg, 2016). We found no effect of recalling positive memories on temporal discounting in older adults, in line with a previous study finding no effect of episodic future thinking on temporal discounting in older adults (Sasse, Peters, & Brassen, 2017). Together, these studies suggest that memory-based manipulations have limited effectiveness in populations with impaired episodic memory. Though there was no overall effect of memory recall on temporal discounting, we did find that the memories that contained more episodic detail (specifically, more event and time details) were more likely to yield patient choice after they were retrieved. No other aspects of the memories, including self-reported ratings of emotional intensity and vividness, were related to patient choice. This further supports the notion that more vivid episodic memory mediates the effect of memory retrieval on choice, using an in-depth autobiographical memory scoring protocol rather than merely self-reported ratings of memory vividness (Peters & Büchel, 2010).

One puzzling aspect of our results is that while place and perceptual details, but not event details, were associated with temporal discounting rates at baseline, event details, but not place or perceptual details, predicted whether memory recall influenced choice. This suggests that the processes contributing to stable time preferences may be distinct from those supporting the flexibility of choice at the time of the decision (Lempert & Phelps, 2016). In line with this idea, individuals with hippocampal damage show similar temporal discounting rates compared to healthy controls (Kwan et al., 2012), but unlike healthy controls their intertemporal choices are not affected by episodic future thinking (Palombo et al., 2015). Perhaps the hippocampus is critical for manipulations that engage episodic retrieval to influence intertemporal choices, while other regions such as the ERC are sufficient for future-oriented choice under normal conditions. Intriguingly, hippocampal volume was most strongly associated with the memory details (time and event) that predicted a change in temporal discounting after autobiographical recall, whereas ERC thickness was associated with the memory details (time, place and perceptual) that predicted temporal discounting at baseline.

In summary, here we show, in a group of cognitively normal older adults, that autobiographical memory richness is correlated with temporal discounting. Time, place, and perceptual details are associated with temporal discounting rates across individuals, and recalling memories that are richer in time and event details prior to choice leads to more patient decisions. These findings will help to inspire and optimize interventions to nudge intertemporal choice, especially in older adults with more limited episodic memory ability. They also add to the growing literature on the critical role of episodic memory in making decisions about the future.

## Supplemental Materials

### Supplemental Methods

#### Self-report questionnaires

After the decision making task was completed, participants filled out four questionnaires on a computer: the Interpersonal Reactivity Index (IRI; Davis, 1983), the Life Orientation Test-Revised (LOT-R; Scheier, Carver, & Bridges, 1994), the Geriatric Depression Scale (GDS; Yesavage, 1988), and the Vividness of Visual Imagery questionnaire (VVIQ; Marks, 1973). The IRI assesses perspective-taking and empathy abilities, abilities that have recently been shown to be associated with temporal discounting (Soutschek, Ruff, Strombach, Kalenscher, & Tobler, 2016). The IRI has four subscales: Perspective-Taking, Empathic Concern, Personal Distress, and Fantasy. We focused analyses on the IRI Perspective-Taking subscale, as this was the one we expected to be most strongly related to temporal discounting based on previous research. The LOT-R tests for optimism, which tends to be elevated in older adults (Mather & Carstensen, 2005), and may be related to future-oriented decision-making (Berndsen & van der Pligt, 2001). The GDS was included as a screening tool, because symptoms of depression are associated with deficits in memory ability, especially in positive memory recall (Dillon, 2015). Therefore, anyone with a GDS score of 9 or above (out of 15), indicating moderate or severe depression, was excluded. Finally, the VVIQ instructs participants to imagine different scenarios in order to measure individual differences in self-reported imagery vividness. VVIQ scores have been shown to be correlated with temporal discounting (Parthasarathi, McConnell, Luery, & Kable, 2017). We examined Spearman correlations between scores on these questionnaires and age, as well as with the size of the effect of the positive memory recall manipulation in our study. Outliers (scores that were more than 2.5 SD from the mean) were removed.

#### List of cues used to elicit positive memory recall on Day 1

Visit from friend/relative from out of town
Participating in sports
Graduating
Birth of a child
Going to the beach
Being in a wedding
Getting engaged or married
Going to a museum
Going to a concert
Going to the theater
Getting a pet
Hosting a party
Favorite team winning a championship
A friend’s birthday party
Fourth of July
Thanksgiving
Winning an award
Getting a job/college/program acceptance letter
Being on a ship/boat
Memorable meal
Class or family reunion
Being promoted/given a raise
Camping or hiking
Buying a house or apartment

#### List of choice sets in the intertemporal choice task (immediate reward amount was always $10)

| Delayed amount (\$) | Delay (days) |
| --- | --- |
| 17 | 1 |
| 20 | 2 |
| 18 | 3 |
| 20 | 4 |
| 24 | 5 |
| 27 | 6 |
| 13 | 7 |
| 14 | 7 |
| 16 | 8 |
| 19 | 9 |
| 25 | 10 |
| 29 | 11 |
| 14 | 13 |
| 25 | 15 |
| 19 | 17 |
| 26 | 18 |
| 16 | 20 |
| 12 | 24 |
| 15 | 26 |
| 32 | 27 |
| 21 | 31 |
| 28 | 38 |
| 15 | 42 |
| 13 | 59 |
| 34 | 60 |
| 35 | 62 |
| 22 | 67 |
| 11 | 74 |
| 18 | 75 |
| 33 | 76 |
| 13 | 86 |
| 12 | 90 |
| 18 | 91 |
| 12 | 96 |
| 15 | 100 |
| 11 | 105 |
| 32 | 107 |
| 11 | 117 |
| 15 | 126 |
| 24 | 127 |
| 14 | 129 |
| 12 | 135 |
| 11 | 137 |
| 12 | 138 |
| 19 | 141 |
| 14 | 154 |
| 25 | 158 |
| 11 | 166 |
| 11 | 167 |
| 19 | 171 |
| 11 | 173 |
| 13 | 176 |
| 12 | 179 |
| 11 | 180 |

### Supplemental Results

#### Self-reported perspective-taking is associated with reduction in discounting following memory recall across participants

The VVIQ, which measures individual differences in self-reported imagery vividness, was uncorrelated with our manipulation’s effect size (N = 33, 1 outlier excluded; ρ = -0.03; p = 0.883) or temporal discounting rate (ρ = -0.09; p = 0.627). However, it was associated with age (ρ = -0.71; p < 0.001), delayed memory recall (ρ = 0.41; p = 0.019), and semantic fluency (ρ = 0.40; p = 0.022) in the expected direction (better memory and younger age are associated with more vivid mental imagery). With respect to autobiographical memory details, VVIQ was associated with event details (ρ = 0.36; p = 0.037), but not perceptual (ρ = 0.27; p = 0.13), emotion/thought (ρ = 0.32; p = 0.069), time (ρ = 0.06; p = 0.72), or place (ρ = -0.002; p = 0.993) details.

The LOT-R, which measures optimism, was associated with the effect size at a trend level (N = 34; ρ = 0.30; p = 0.088), as well as with delayed memory recall (ρ = 0.47; p = 0.005) and semantic fluency (ρ = 0.54; p = 0.001). People who are more optimistic about the future tend to have better memory ability. LOT-R scores were not associated with age (ρ = -0.06; p = 0.721), Control condition temporal discounting rate (ρ = 0.03; p = 0.884), or any autobiographical memory details (event: ρ = 0.14, p = 0.417; time: ρ = 0.29, p = 0.095; place: ρ = 0.19; p = 0.293; emotion: ρ = -0.08; p = 0.639; perceptual: ρ = 0.01; p = 0.934).

The IRI perspective-taking subscale measures an individual’s tendency to consider the perspective of others in everyday situations. There was a significant association between an individual’s perspective-taking score and the effect size of the manipulation (N = 33; ρ = 0.60; p < 0.001; Fig. 4), in that individuals with a greater propensity and/or capacity to take the perspective of others were more likely to show reduced temporal discounting after recalling positive memories. This association was robust to controlling for the covariates of age, gender, and years of education (partial ρ = 0.55; p = 0.002). Perspective-taking was unrelated to temporal discounting rate in the Control condition (ρ = 0.14; p = 0.453), delayed memory recall (ρ = 0.15; p = 0.397), semantic fluency (ρ = 0.23; p = 0.190), and number of autobiographical memory details (event details: ρ = 0.13, p = 0.483; time: ρ = 0.15, p = 0.403; place: ρ = -0.16; p = 0.368; emotion: ρ = 0.11; p = 0.542; perceptual: ρ = 0.18; p = 0.308).

#### Participant ratings of memories did not predict change in choice following recall of memories

We investigated whether any characteristics of the memories themselves, as rated by the participants, could predict the extent to which retrieving them was effective in reducing temporal discounting rate. We conducted a series of mixed-effects logistic regressions to see which ratings could predict choice of delayed reward on a trial-by-trial basis, controlling for the subjective value of rewards computed assuming the Control condition discount rate. None of the ratings were significant predictors of choice: the Day 1 rating of “similarity between past and present self” (Coefficient = -0.028; p = 0.792), the Day 1 rating of “feeling when recalling the memory now” (Coefficient = -0.092; p = 0.588), the Day 1 rating of “emotional intensity” (Coefficient = - 0.184; p = 0.179), the Day 1 rating of “feeling during the memory” (Coefficient = -0.330; p = 0.056), the Day 1 rating of “personal importance of the memory” (Coefficient = -0.087; p = 0.470), the Day 1 rating of memory vividness (Coefficient = 0.118; p = 0.476), the Day 2 rating of valence (Coefficient = 0.099; p = 0.893), the Day 2 rating of emotional intensity (Coefficient = 0.125; p = 0.410), and the Day 2 rating of “feeling when recalling the memory now” (Coefficient = -0.048; p = 0.771) all yielded null results.

**Supplemental Fig. 1.**
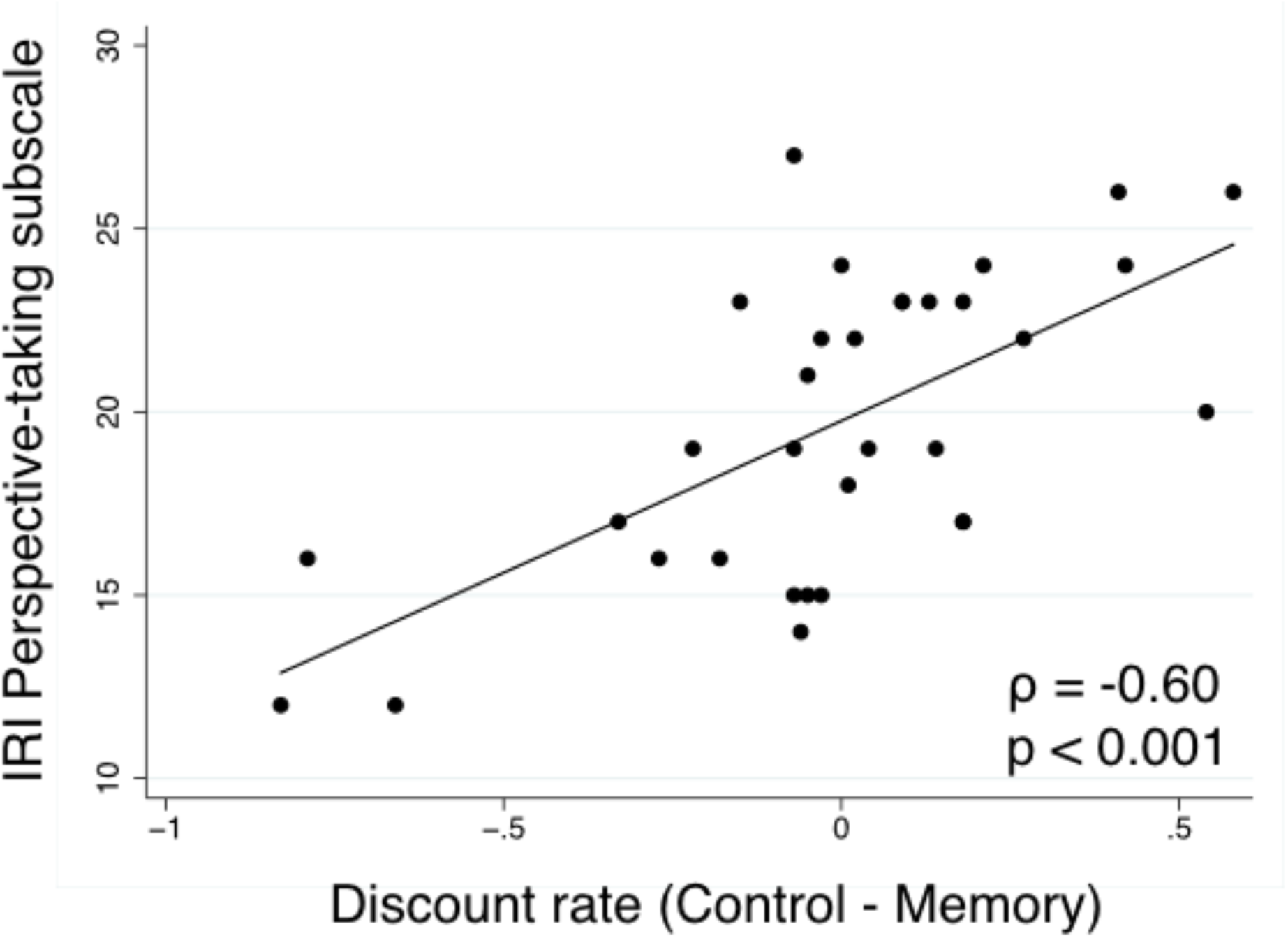
The perspective-taking subscale of the Interpersonal Reactivity Index was significantly associated with the degree to which discount rates were reduced after participants recalled positive memories.

**Supplemental Table 1.**
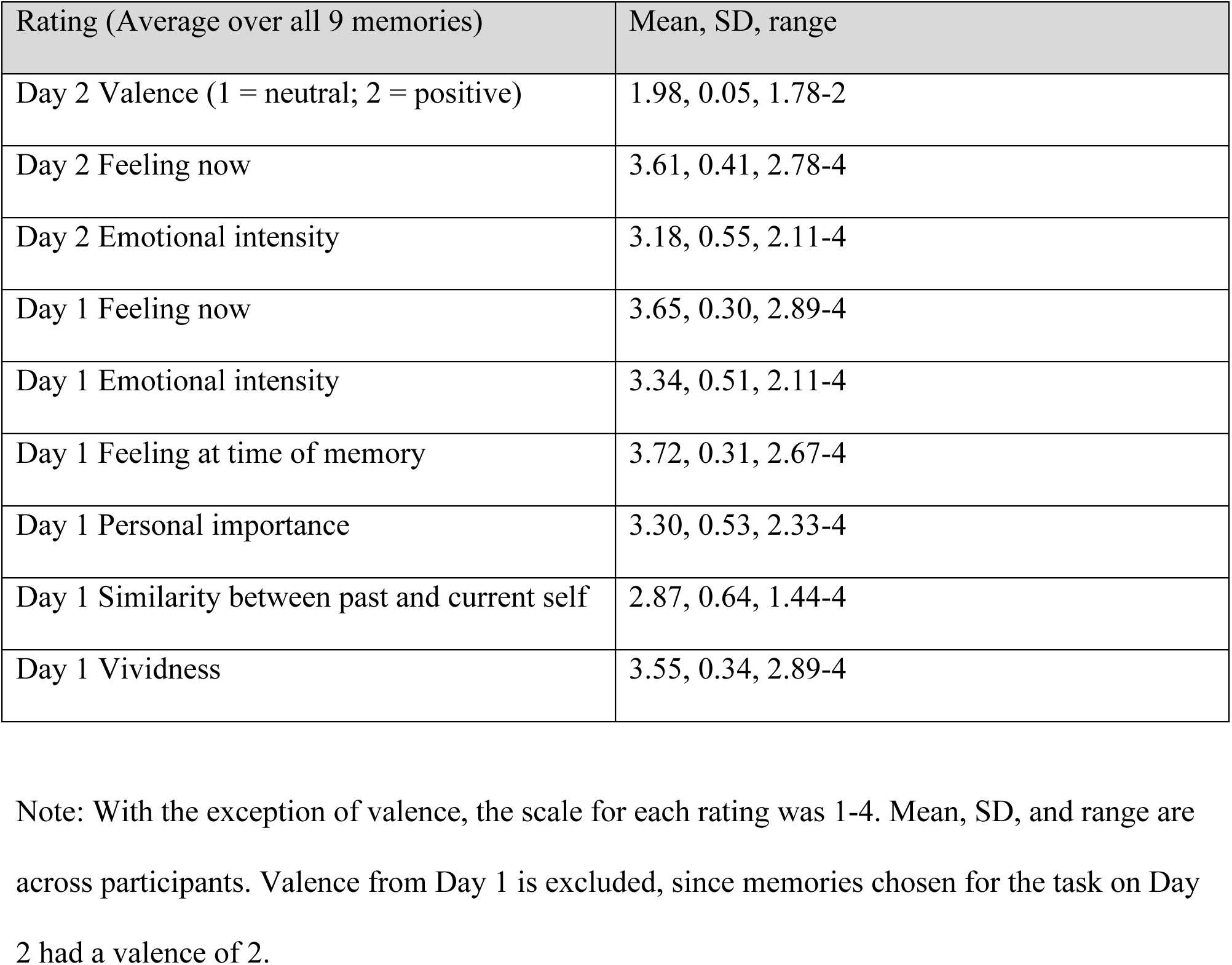
Average ratings of memories recalled during intertemporal choice task.

| Rating (Average over all 9 memories) | Mean, SD, range |
| --- | --- |
| Day 2 Valence (1 = neutral; 2 = positive) | 1.98, 0.05, 1.78-2 |
| Day 2 Feeling now | 3.61, 0.41, 2.78-4 |
| Day 2 Emotional intensity | 3.18, 0.55, 2.11-4 |
| Day 1 Feeling now | 3.65, 0.30, 2.89-4 |
| Day 1 Emotional intensity | 3.34, 0.51, 2.11-4 |
| Day 1 Feeling at time of memory | 3.72, 0.31, 2.67-4 |
| Day 1 Personal importance | 3.30, 0.53, 2.33-4 |
| Day 1 Similarity between past and current self | 2.87, 0.64, 1.44-4 |
| Day 1 Vividness | 3.55, 0.34, 2.89-4 |

Note: With the exception of valence, the scale for each rating was 1-4. Mean, SD, and range are across participants. Valence from Day 1 is excluded, since memories chosen for the task on Day 2 had a valence of 2.

